# Spatiotemporal patterns of desiccation tolerance in natural populations of *Drosophila melanogaster*

**DOI:** 10.1101/079616

**Authors:** Subhash Rajpurohit, Eran Gefen, Alan Bergland, Dmitri Petrov, Allen G Gibbs, Paul S Schmidt

## Abstract

Water availability is a major environmental challenge to a variety of terrestrial organisms. In insects, desiccation tolerance varies predictably over various spatial and temporal scales and is an important physiological basis of fitness variation among natural populations. Here, we examine the dynamics of desiccation tolerance in North American populations of *Drosophila melanogaster* using: 1) natural populations sampled across latitudes and seasons in the eastern USA; 2) experimental evolution in the field in response to changing seasonal environments; 3) a sequenced panel of inbred lines (DGRP) to perform genome wide associations and examine whether SNPs/genes associated with variation in desiccation tolerance exhibit patterns of clinal and/or seasonal enrichment in pooled sequencing of populations. In natural populations we observed a shallow cline in desiccation tolerance, for which tolerance exhibited a positive association with latitude; the steepness of this cline increased with decreasing culture temperature, demonstrating a significant degree of thermal plasticity. No differences in desiccation tolerance were observed between spring and autumn collections from three mid-to-northern latitude populations, or as a function of experimental evolution to seasonality. Similarly, water loss rates did not vary significantly among latitudinal, seasonal or experimental evolution populations. However, changes in metabolic rates during prolonged exposure to dry conditions indicate increased tolerance in higher latitude populations. Genome wide association studies identified thirty-six SNPs in twenty-eight genes associated with sex-averaged drought tolerance. Among North American populations, genes associated with drought tolerance do not show increased signatures of spatially varying selection relative to the rest of the genome, whereas among Australian populations they do.

## Introduction

Insects exploit and inhabit a wide range of habitats on planet earth which range from hot deserts to cold arctic regions. In many of these environments presence of water is scarce and desiccation is a major threat to many terrestrial organisms living there. Insects are particularly most vulnerable to water related challenges, because of their small size and thus large surface area to volume ratio. (Gibbs & Rajpurohit 2010). Environmental stresses such as desiccation are highly variable among these natural habitats, and often vary predictably with such features as latitude, altitude, and season. Patterns of phenotypic and genetic variation distributed along these gradients, both within and among taxa, offer means to address the extent to which divergence in physiologically selected traits are affected by natural selection (Barton 1999; Whitlock & McCauley 1999).

*Drosophila* species constitute good models for studies relating to population ecology and physiological adaptations (Parsons 1983; Lemeunier et al. 1986). During their evolutionary history, different *Drosophila* species have adapted to diverse climatic conditions. This has been clearly demonstrated on multiple continents where multiple Drosophila species exhibit pronounced clines for multiple fitness traits (e.g., Hoffmann & Parsons 1991; Hoffmann & Parsons 1993; Blows & Hoffmann 1993; Karan et al. 1998; Gilchrist et al. 2004; Schmidt et al. 2005; Wittkopp et al. 2011; Rajpurohit & Nedved 2013). In various taxa, high levels of desiccation resistance are associated with adaptation to arid habitats (e.g., David et al. 1983; Hoffmann and Parsons 1991; Gibbs et al. 2003). Several studies have reported geographical variation for desiccation tolerance in *Drosophila* species inhabiting arid environments characterized by elevated temperatures (e.g., Hoffmann & Harshman 1999; Matzkin et al. 2007; Rajpurohit & Nedved 2013; Rajpurohit et al. 2013b).

In the cosmopolitan *D. melanogaster*, latitudinal clines have been observed for a wide range of fitness-associated traits including thermal tolerance, body size, basic life histories, incidence of reproductive dormancy, cuticular hydrocarbon composition, and aspects of behavior (e.g., David 1975; Coyne and Beecham 1987; James et al. 1995, 1997; Zwaan et al. 2000; Schmidt et al. 2005; De Moed et al. 1997; Mitrovski and Hoffmann 2001; Karan et al. 1998; Eanes 1999; Verrelli and Eanes 2001; Lemeunier et al. 1986; Rajpurohit & Schmidt 2016). While such patterns may be influenced by demography (Kao et al. 2015; Bergland et al. 2016), latitudinal clines in *D. melanogaster* are commonly interpreted as resulting from spatially varying selection and local adaptation to climatic and associated variables.

However, the role of desiccation tolerance in the adaptation of *D. melanogaster* to environmental heterogeneity has not been comprehensively addressed (Hoffmann et al. 2001; Hoffman et al. 2005; Telonis-Scott et al. 2006; Parkash et al. 2008); there has been little work done on *D. melanogaster* populations inhabiting temperate latitudinal ranges, in which humidity and associated desiccation stress are predicted to vary both with latitude and season. Similarly, the genetic basis of variation in desiccation tolerance remains unresolved (but see Telonis-Scott et al. 2016). We therefore carried out an integrated study on desiccation tolerance in natural and experimental populations in a genetic model organism, utilizing a combination of phenotypic, physiological and plasticity analyses, population level sequencing and genome wide association studies. Our results demonstrate thermally mediated plasticity, associations among latitude, physiology and stress tolerance, and identify a set of candidate SNPs that may underlie variation in desiccation tolerance in natural populations.

## Material & Methods

### Fly collection and maintenance

Six natural populations of *D. melanogaster* were collected from fruit orchards located along the east coast of the United States by a combination of aspiration and baiting/sweeping with aerial nets (see Fig. 1). Gravid females were immediately sorted into isofemale lines in the field; once the resulting progeny eclosed, lines were typed to species. Approximately 150 isofemale *D. melanogaster* lines were collected from each locale. Populations sampled were: Bowdoin, Maine (44.03N, 73.20W); Media, Pennsylvania (40.04N, 76.30W); Charlottesville, Virginia (38.03N, 78.48W); Athens, Georgia (32.84N, 83.66W); Jacksonville, Florida (30.33N, 81.66W); and Homestead, Florida (25.46N, 80.45W). Seasonal collections were done in Media, Pennsylvania, Lancaster, Massachusetts (42.455N, 71.67W) and Charlottesville, Virginia (38.03N, 78.48W) orchards in June and October of 2012. Long-term maintenance of all populations was done at 24 °C, 12:12 light:dark photoperiod, with a generation time of approximately 21 days. Flies were maintained on regular cornmeal-molasses-agar media.

**Fig. 1:**
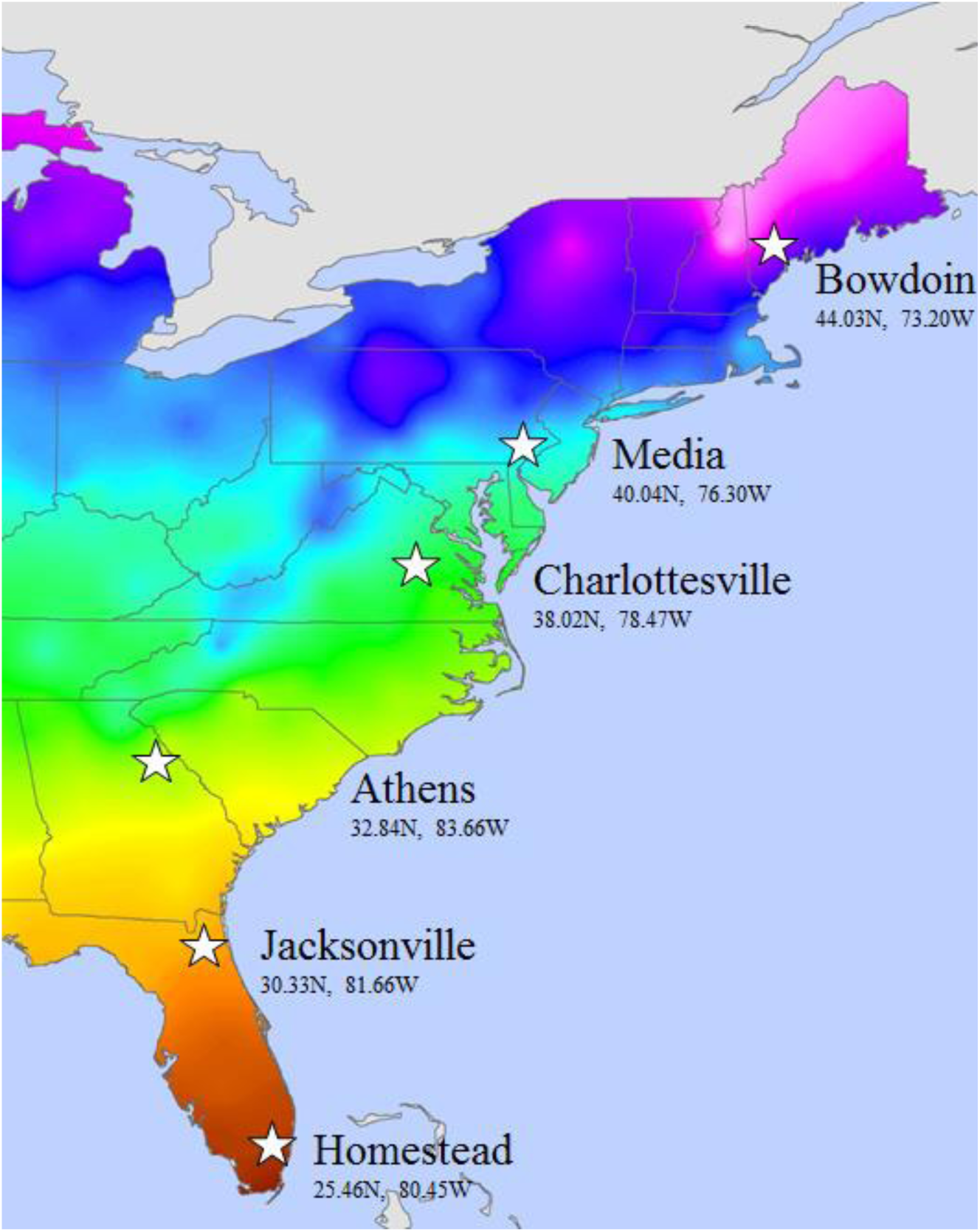
Temperature map of the east coast of the U.S.A. showing the populations of *D. melanogaster* that were collected and assayed. Seasonal collections were done in Lancaster, MA, Media, PA and Charlottesville, VA in 2012.

### Establishment of clinal cages

To establish population cages for each spatial and temporal collection, we pooled 25 isofemale lines. We created two population cages for each of the spatial and temporal collections (see above). Each cage was created using independent sets of 25 isofemale lines by releasing ten mated females from each line into 12x12x12 inch insect enclosures (Live Monarch Foundation, Boca Raton, Florida, USA). These lines were maintained in mass culture and allowed to outcross for 5 generations; subsequently, samples were collected for the phenotypic assays described below.

### Desiccation tolerance assay

Four-to-five day old virgin flies were transferred to empty vials in groups of ten, and restricted to the lower half of the vials by a foam stopper. Silica gel was then added above the stopper to maintain low humidity, and the vial was sealed with Parafilm^TM^. Mortality was recorded at hourly intervals until all flies were dead. Fifteen to twenty-five isofemale lines per population were assayed. Throughout the desiccation tolerance assay vials were kept in a 25 °C incubator.

### Thermal plasticity and diapause treatment

To examine thermal plasticity for desiccation tolerance in the latitudinal collections, we examined tolerance as a function of 1) three culture temperatures (18, 25 and 28 °C), and 2) after exposure to conditions that elicit reproductive dormancy (3-week exposure to 11˚C, 10L:14D photoperiod). Each replicate population cage for each of the geographic regions was allowed to oviposit for a 2-3h at 25 °C in successive culture vials; egg density was standardized at 40-50 eggs per vial by manual removal. Replicate vials were then randomly assigned to one of the four experimental treatments and cultured in Percival I36VL incubators. On emergence, adults were transferred to fresh food vials and aged for 4-5 days before subjecting them to the desiccation tolerance assay; for the reproductive dormancy assay, flies were immediately evaluated for desiccation tolerance following the 3-week period of exposure.

### Experimental manipulation of desiccation tolerance evolution

To examine how desiccation tolerance evolves under field conditions, we established 20 experimental mesocosms at a field site in Philadelphia, PA USA. Each mesocosm was an 8m^3^ outdoor insect rearing enclosure (Bioquip Products, Gardena, CA) surrounding a mature (dwarf) peach tree. Each mesocosm was seeded with 1000 individuals (500 males, 500 females) derived from a collection made in 2012 from the same PA orchard as described above. This progenitor population was created by pooling 86 independent inbred lines, allowing them to recombine and expand for 10 generations in the laboratory, and then maintained at large census size (~10^6^) in the laboratory for the duration of the experiment (July 1 – November 1, 2014). Each cage was randomly assigned to a treatment: seasonally evolving (E) or non-seasonally evolving (F). In the E treatment, populations were supplied with fresh food/oviposition sources (500ml standard cornmeal-molasses medium) every 2d, and the populations were allowed to evolve and adapt to seasonal environmental conditions. In the F treatment, fresh food/oviposition sources were also supplied: however, all eggs laid by the experimental flies were counted, removed, and replaced with the same number of eggs from the progenitor laboratory population that was maintained under aseasonal, laboratory conditions. Thus, the F populations were maintained under normal demographic trajectories and flies were directly exposed to the field environment, but these populations were not allowed to evolve to the field conditions; the F treatment represents the progenitor laboratory population while including any potential epigenetic marks that may be elicited upon exposure to field conditions during direct development as well as any evolution in the aseasonal, laboratory environment. At the end of the experiment, a sample of approximately 2000 eggs was collected from each of the 20 cages and brought back to the laboratory. These collections were allowed to develop in the laboratory and subsequently passed through two generations of common garden, density controlled culture. In the F3 generation subsequent to the field collection, experimental animals were collected and processed for respirometry measurements.

### Respirometry measurements

Within 3 hours of adult eclosion, virgin males and females were collected, and sets of fifteen were placed in fresh food vials. A total of 24 sets were assayed for the latitudinal extremes of Maine and Florida (2 geographic regions × 2 sexes × 2 replicate populations × 3 experimental replicates), 24 for seasonal comparisons in the Pennsylvania orchard (2 seasons × 2 sexes × 2 replicate cages × 3 repeats) and 40 for experimental evolution (2 treatments × 2 sexes × 10 replicate cages). Respirometry was carried out at 4-6 days post-eclosion on sets of 10-15 individuals. Flies were transferred directly from their food vials to a 4 mL glass metabolic chamber with aluminum stoppers, which was covered with a black cardboard sleeve to reduce activity in the chamber. Flow through respirometry at 25°C was carried out using two channels of a flow multiplexer (RM-8, Sable Systems International, Las Vegas, NV, USA), where dry, CO_2_-free air was supplied to the chambers at 50 mL·min^−1^ using factory-calibrated mass flow controllers (MC-500 sccm; Alicat Scientific, Tuscon, AZ, USA), and excurrent air from the measured chamber was passed through a LI-7000 CO_2_/water vapor dual analyzer (Li-Cor Biosciences, Lincoln NE, USA). The flies were acclimated to the experimental chambers and air flow for 15 minutes during which flies from the alternative chamber were measured. Identical empty chambers were used for baselining. Analyzer voltage output was recorded, stored and analyzed using UI-2 data acquisition interphase and Expedata software (Sable Systems International). Recording rate was set to 1 Hz and only data from the last 5 minutes of each 15 min run were averaged for analysis.

We used an additional experimental approach to compare the Florida and Maine populations for which significant differences in desiccation resistance were found (see results). It was previously reported that water loss rates of *D. melanogaster* under similar experimental conditions stabilize only after >2h of exposure (Gibbs et al. 1997), and therefore we carried out respirometry on additional sets of flies (2 geographic regions × 2 sexes × 2 replicate populations × 6 repeats), which were randomly allocated to six multiplexer channels, with a seventh used for baselining. The measurement sequence of 20 min measurements was as follows: baselining, 3 experimental chambers, baselining, three additional experimental chambers and finally baselining again for a total file recording of 3 h, which was immediately followed by another recording at the same sequence. As each set of flies was added to the respirometry setup 20 min prior to the initial measurement, and only data from the last 10 minutes were analyzed, the flies were assayed 30 min and following additional 3h during which the flies were exposed to experimental temperature and dry air flow conditions.

### Desiccation assays for the Drosophila Genetic Reference Panel (DGRP)

DGRP lines (Mackay et al. 2012) were obtained from the Bloomington Drosophila Stock Center and maintained in the lab on yeast-cornmeal-sucrose medium at 25^◦^C. For assays, 4-6 day old flies were sorted by sex and held on fresh media for two days. On the days experiments began, flies were lightly anesthetized with CO_2_ and ten groups of five flies each were rapidly transferred to empty shell vials. A foam plug was inserted halfway down the vial, silica gel desiccant was added to the upper half of the vial, and the vial was sealed with Parafilm^TM^. The vials were transferred to a 25^◦^C incubator with constant illumination, and dead flies were counted hourly until all flies were dead.

Desiccation assays were performed in blocks of ~30 lines per block. Because of the time required to initiate desiccation stress for all lines and to count dead flies, we recorded the exact time when desiccant was added to assay vials and the exact times when each line was checked. Flies that died before the first survival check were assumed to have been injured by handling or other stress and were not included in the data analysis. Any flies dying between two checks were assumed to have died midway between them. To assess potential variation among blocks associated with minor differences in food, incubator temperature, etc., two lines (RAL-315 and RAL-324) were assayed in each block as internal controls. Variation among blocks in desiccation resistance of each line was <10%. For these lines, only data from the first block were included in the overall data analysis.

### Genome wide association analysis

We performed genome-wide association analysis on desiccation tolerance using the Drosophila Genetic Reference Panel (DGRP). Phenotypic line means were uploaded to the DGRP analysis website (http://dgrp2.gnets.ncsu.edu) for genome-wide association analysis following methods outlined in Mackay et al. (2012) and Huang et al. (2014). Thirty-six (36) SNPs were associated with sex-averaged desiccation tolerance below nominal *p-* value of 1e-5 (Fig. S1). These SNPs were annotated as being in or nearby 36 genes (Table S1, Table S2, and Table S3).

We tested whether these 36 genes were more likely to show signals of spatially varying selection than expected relative to the rest of the genome. We examined patterns of spatially varying selection using whole genome resequencing of populations sampled along the east coasts of North America (Bergland et al. 2014) and Australia (Bergland et al. 2016) following a method outlined in Daub et al. (2013). This method tests whether gene sets identified *a priori* show stronger signals of spatially varying selection than sets of control genes. To perform this analysis, we first estimated genetic differentiation at approximately 500,000 common SNPs with average minor allele frequency greater than 5% (Bergland et al. 2014) among populations of *Drosophila melanogaster* along latitudinal transects in North America or Australia using the *T*_*FLK*_ statistic (Bonhomme et al. 2010; Bergland et al. 2016). The *T*_*FLK*_ statistic is a modified version of the classic Lewontin-Krakauer test for *F*_*ST*_ outliers that incorporates certain aspects of population structure and has been shown to have a low false positive rate when sampled populations result from secondary contact, as is likely the case for North American and Australian populations of *D. melanogaster* (Caracristi & Schlotterer 2003; Duchen et al. 2013; Kao et al. 2015; Bergland et al. 2016). *T*_*FLK*_ values were *z-*transformed and, following Daub et al. (2013), we refer to these transformed *T*_*FLK*_ values as *z*. For each gene in the genome, we calculated the maximum *z* value, *z(g)*, by considering all SNPs within 10Kb of the beginning and end of the gene. *z(g)* was normalized by gene length (hereafter *z*_*st*_*(g)*) by binning all genes with approximately equal length on a log_2_ scale following equations (1) and (2) of Daub et al. (2013). Next, we generated 1000 sets of control genes matched to the target set associated with desiccation tolerance. These control sets were matched by chromosome, inversion status at the large cosmopolitan inversions that segregate on each chromosome (Corbett-Detig & Hartl 2012), recombination rate (Comeron et al. 2012), and SNP density per basepair. For target and control gene sets we calculated the sum of *z*_*st*_(*g*), *SUMSTAT*, and estimated the probability that *SUMSTAT*_*target*_ is greater than *SUMSTAT*_*control*_.

### Other statistics

Desiccation tolerance is presented as means ± s.e., and data were analyzed using ANOVA to model main effects and interaction terms. As the investigated populations encounter different temperature and humidity conditions in their natural habitats we performed a multiple regression analysis of trait values as a simultaneous function of T_min_, T_max_, T_ave_, RH_min_, RH_max_ and RH_ave_ of origin of populations. Respirometry data were analyzed with ANOVA when body mass did not vary significantly between experimental groups. On the rare occasion (see results) that body mass did vary, data were analyzed using ANCOVA with body mass as a covariate. JMP v12 (SAS Institute, Cary, NC) and Statistica (Statsoft Inc., Release 7.0, Tulsa, OK, USA) were used for calculations as well as illustrations.

## Results

### Spatiotemporal variation in desiccation tolerance

Populations from six latitudinal localities of *D. melanogaster* (Fig. 1) exhibited a positive cline for desiccation tolerance in both sexes, and desiccation tolerance increased positively with latitude (Fig 2ABC). These patterns were affected by culture temperature, demonstrating an inverse relationship between temperature and desiccation tolerance (Table 1). The significant interaction terms of population by temperature and temperature by sex showed that patterns of thermal plasticity varied among populations and were distinct between the sexes (Table 1; Fig. 2ABC).

**Fig. 2:**
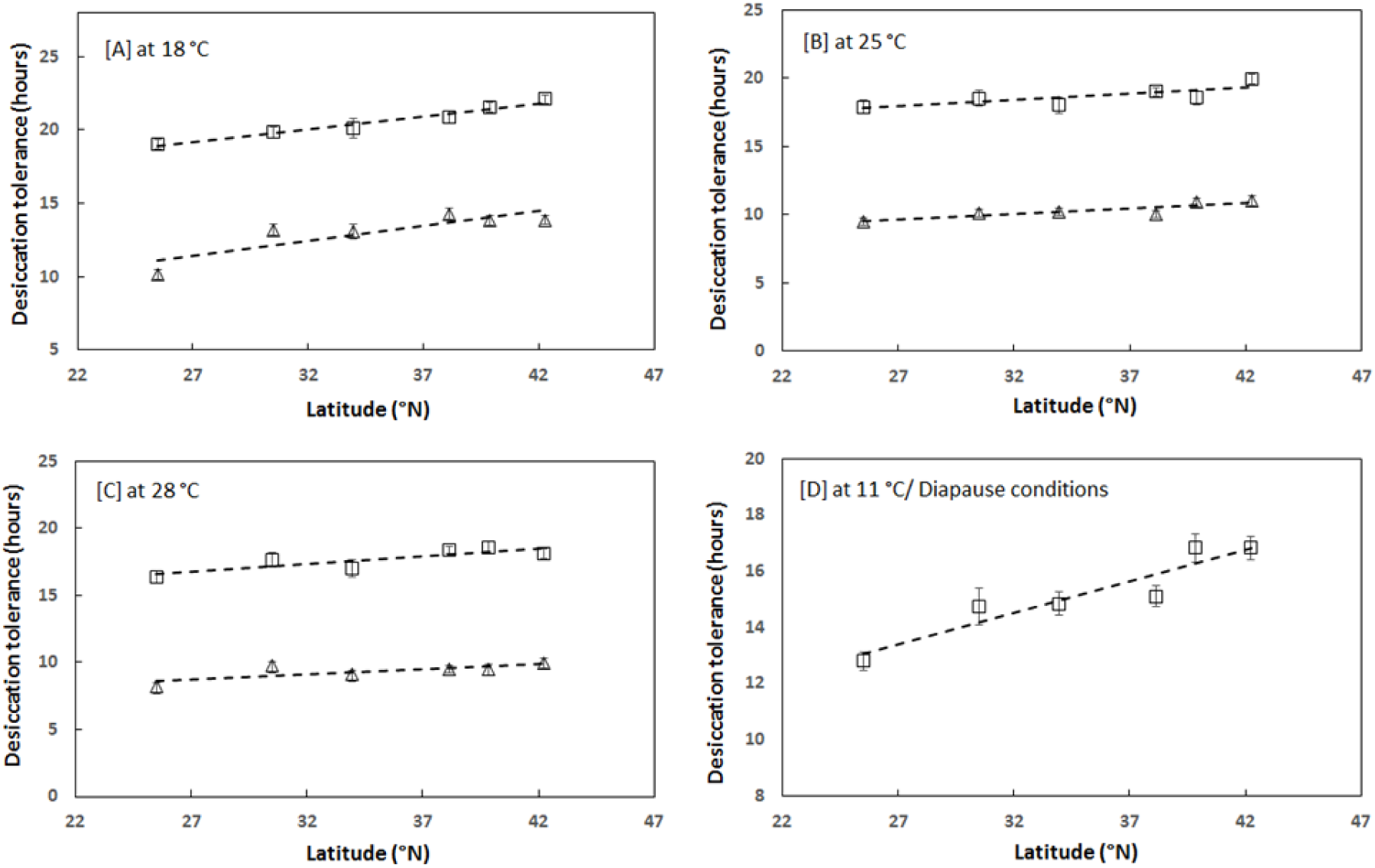
Data (mean±s.e.) on desiccation tolerance for males and females of *D. melanogaster* from six geographical locations (see Fig. 1) at four different thermal conditions (18 °C (A), 25 °C (B), 28 °C (C), and Diapause conditions)))). To study over-wintering affects (D) over desiccation tolerance flies were kept under dormancy inducing conditions (11 °C and 09:15 photoperiod) for 3 weeks before subjecting them to desiccation tolerance assay (at 25 °C). Males and females are denoted as triangles and rectangles respectively.

**Table 1:** ANOVA for desiccation tolerance of males and females from the six geographically distinct populations.

| **Parameters** | **SS** | **d. f.** | **MS** | **F** | ***p*** |
| --- | --- | --- | --- | --- | --- |
| Population (P) | 532.2 | 5 | 106.4 | 28.74 | **0.000000** |
| Temperature (T) | 1709.3 | 2 | 854.7 | 230.75 | **0.000000** |
| Sex (S) | 14039.0 | 1 | 14039.0 | 3790.42 | **0.000000** |
| P x T | 104.6 | 10 | 10.5 | 2.82 | **0.001866** |
| P x S | 27.5 | 5 | 5.5 | 1.49 | 0.191275 |
| T x S | 33.1 | 2 | 16.6 | 4.47 | **0.011721** |
| P x T x S | 64.3 | 10 | 6.4 | 1.74 | 0.068731 |
| Error | 3229.7 | 872 | 3.7 |  |  |

Similarly, we observed a strong, positive correlation between desiccation tolerance and latitude in adult females that had been exposed for three weeks to environmental conditions that induce reproductive quiescence (Fig. 2D). This demonstrates that physiological plasticity at the adult life stage also impacts desiccation tolerance. Furthermore, the differences among populations were maximized when assayed as adults following this three-week exposure, although this may also be related to geographic differences in the incidence of reproductive dormancy (Schmidt et al. 2005). The overall duration of survivorship in the desiccation assay was lower for older, post-dormancy than in young adults, which could simply reflect biological age.

Regression analysis of desiccation tolerance with geographical and climatic variables (T_min_, T_max_, T_ave_, RH_min_, RH_max_ and RH_ave_) for both sexes at all three culture temperatures is given in Table 2. Temperature parameters exhibited stronger associations with desiccation tolerance than did humidity parameters. Temperature emerged more significant than humidity parameters for the variations observed in desiccation tolerance along the east coast of USA. Interestingly, minimum temperature displayed the most robust association with desiccation tolerance in the geographic collections. This is consistent with the hypothesis that desiccation tolerance may have stronger effects on performance and fitness during cooler periods in temperate environments.

**Table 2:** Regression analysis of desiccation tolerance with geographical and climatic parameters. The first row gives correlations (r values), the second line the goodness of fit from ANOVA. Statistical significance is depicted by bold typeface.

| Temperature |  | Latitude | T<sub>min</sub> | T<sub>max</sub> | T<sub>ave</sub> | RH<sub>min</sub> | RH<sub>max</sub> | RH<sub>ave</sub> |
| --- | --- | --- | --- | --- | --- | --- | --- | --- |
| 18 °C Male | r | 0.8582 | -0.8766 | -0.6603 | -0.7901 | -0.9236 | -0.1267 | -0.6808 |
|  | *p* | **0.02874** | **0.0219** | 0.15353 | 0.06148 | **0.00854** | 0.81104 | 0.13654 |
| 18 °C Female | r | 0.98115 | -0.9531 | -0.9651 | -0.977 | -0.6814 | -0.5908 | -0.7623 |
|  | *p* | **0.00053** | **0.00325** | **0.0018** | **0.00079** | 0.13612 | 0.2169 | 0.07804 |
| 25 °C Male | r | 0.87516 | -0.8453 | -0.8305 | -0.8544 | -0.6351 | -0.4225 | -0.5673 |
|  | *p* | **0.02241** | **0.03405** | **0.04064** | **0.03028** | 0.17541 | 0.40394 | 0.24034 |
| 25 °C Female | r | 0.78074 | -0.82 | -0.8713 | -0.8602 | -0.3712 | -0.3493 | -0.4461 |
|  | *p* | 0.06684 | **0.04567** | **0.02377** | **0.02797** | 0.46884 | 0.4974 | 0.37529 |
| 29 °C Male | r | 0.76591 | -0.8095 | -0.6357 | -0.742 | -0.7053 | 0.06338 | -0.3727 |
|  | *p* | 0.07579 | 0.05098 | 0.17491 | 0.09126 | 0.11748 | 0.90505 | 0.46687 |
| 29 °C Female | r | 0.86563 | -0.8094 | -0.7261 | -0.7852 | -0.7537 | -0.3719 | -0.7134 |
|  | *p* | **0.02587** | 0.05102 | 0.10226 | 0.06426 | 0.08353 | 0.46788 | 0.11146 |

Seasonal variation in desiccation tolerance for the three sampled populations (MA, PA, and VA) is depicted in Figure 3. Again, we observed significant variation in desiccation tolerance among populations, and the significant three-way interaction term for population x season x sex indicated that the change in desiccation tolerance from spring to fall varied by sex and population. Contrary to our predictions, however, we did not observe any consistent difference in desiccation tolerance between spring and fall samples (Table 3; F=3.01; d.f.=1; *p*=0.08).

**Fig. 3:**
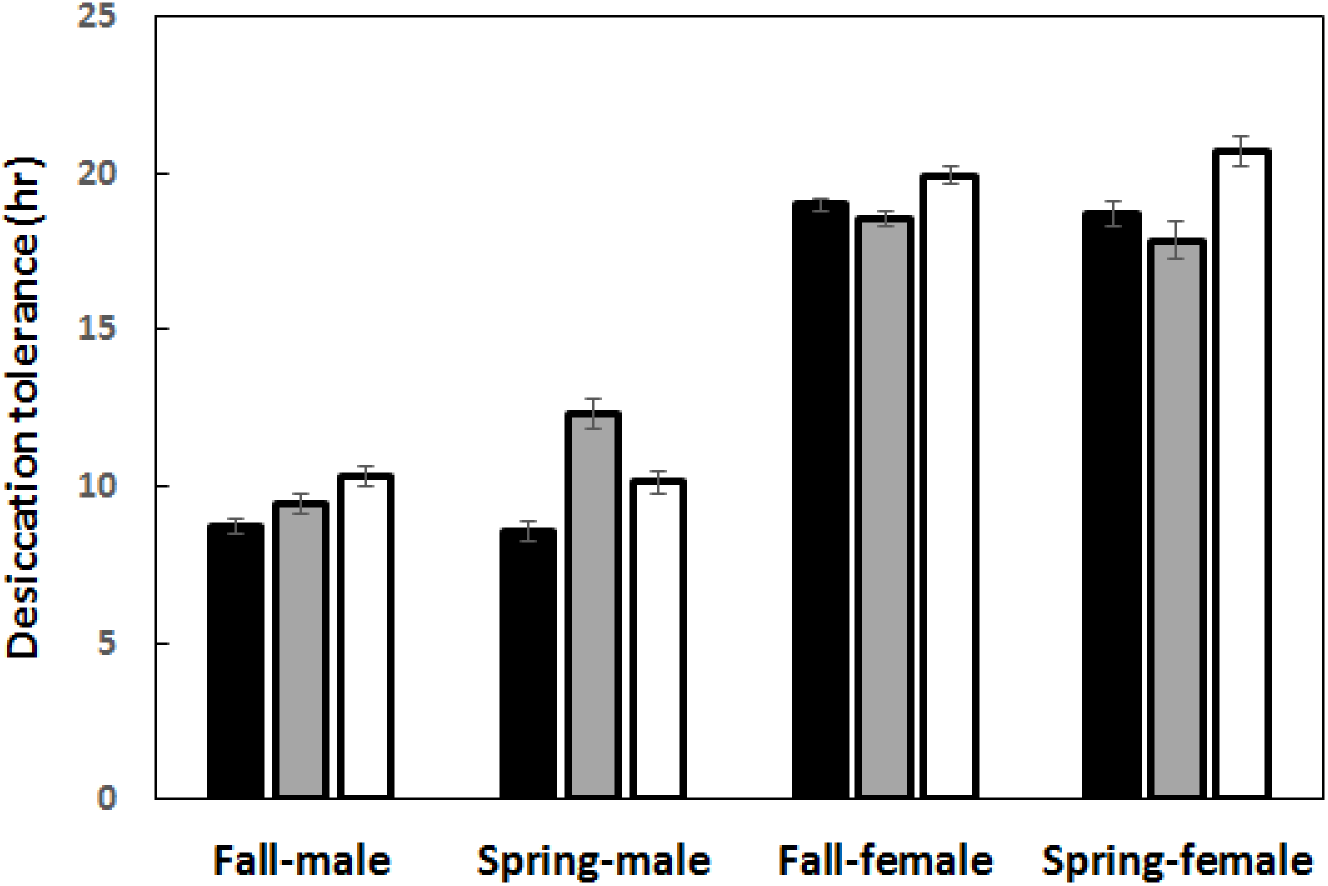
Data (mean±s.e.) on early and late season desiccation tolerance for males and females of *D. melanogaster* from three geographical locations (Lancaster, MA; Media, PA; Charlottesville, VA). Open, gray and black bars represent MA, PA and VA populations respectively.

**Table 3:** ANOVA for seasonal variations in desiccation tolerance of males and females from the three geographically distinct populations.

| **Parameters** | **SS** | **d.f.** | **MS** | **F** | ***p*** |
| --- | --- | --- | --- | --- | --- |
| Population (Pop) | 109.66 | 2 | 54.83 | 14.95 | **0.000001** |
| Season (Sea) | 11.06 | 1 | 11.06 | 3.01 | 0.083603 |
| Sex | 6188.14 | 1 | 6188.14 | 1686.67 | **0.000000** |
| Pop*Sea | 21.49 | 2 | 10.74 | 2.93 | 0.055111 |
| Pop*Sex | 133.73 | 2 | 66.87 | 18.23 | **0.000000** |
| Sea*Sex | 14.78 | 1 | 14.78 | 4.03 | **0.045685** |
| Pop*Sea*Sex | 67.72 | 2 | 33.86 | 9.23 | **0.000131** |
| Error | 1034.62 | 282 | 3.67 |  |  |

### Water loss rate and metabolic rate in natural and experimental populations

With the exception of Florida and Maine males, we did not find a significant within-sex difference in mean body mass among all our comparisons (data not shown). The mean body mass of male and female flies (ca. 0.7-0.8 and 1.1-1.2 mg, respectively) correlated with the well-documented higher desiccation resistance of females (e.g. Gibbs et al. 1997). We found no significant difference for either sex in mean 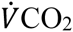 or water loss rates (WLRs) between Florida and Maine, early and late season collections from Pennsylvania, or evolving and fixed populations (ANOVA, with replication as a random factor, p>0.05). When comparing Florida and Maine males, no significant difference was found in mean 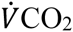 (F_1,7_=1.61, p=0.24) or water loss rates (F_1,7_=2.76, p=0.21), even when accounting for body size (ANCOVA, with body mass as a covariate).

The limitation of our original respirometry approach in detecting variation in WLRs prompted us to focus on the two most geographically-distant populations for which significant variation in desiccation resistance were detected (i.e. Florida vs. Maine; see Fig. 2), using an alternative approach (see Methods). Still, no significant difference was found in the WLRs among female (ANOVA; F_1,21_=0.005, p=0.95) or male (F_1,21_=0.13, p=0.71) flies even when exposed to dry air flow for 3 h. No significant differences were found in 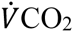 between Florida and Maine females (p=0.68 and 0.34 after 30 min and 3 h, respectively), but similar values for males after 30 minutes (p=0.46) were followed by significantly lower 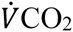 values for Maine compared with Florida males (F_1,21_=4.34, p=0.049). Interestingly, when testing for temporal changes within sets of flies we found a significant increase in 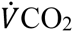 of Florida males from initial values (3.16±0.15 µL·fly^−1^·h^−1^) compared to those recorded after 3 h of exposure to desiccation (3.82±0.13 µL·fly-1·h-1) (paired t-test; t_11_=5.04, p<0.001). In contrast, values after 30 min and 3 h of desiccation did not vary significantly for Maine male flies (t11=1.89, p=0.09) (Fig. 4A). Among female flies, a significant decrease in 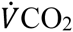 with exposure to desiccation was recorded for Maine (t_11_=3.69, p=0.004), but not Florida populations (t_11_=0.40, p=0.70) (Fig. 4B).

**Fig. 4:**
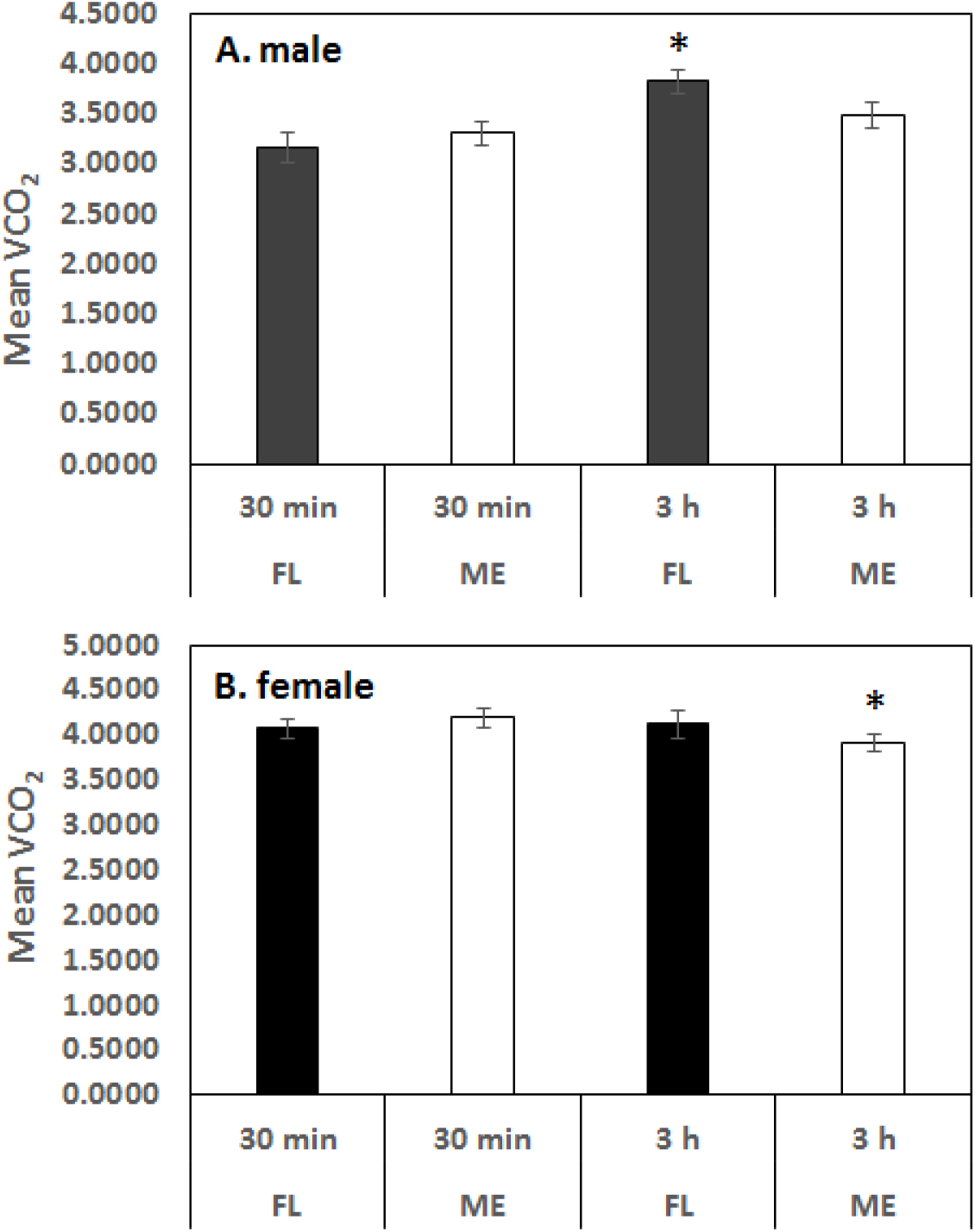
Carbon dioxide emission rates for males (A) and females (B) of *D. melanogaster* populations from Florida (FL) and Maine (MN) locations. Measurements were done at two time points (30 min and 3hr). Asterisks indicate significant difference from 30 min values (α=0.05).

### GWAS, enrichment and parallelism of SNPs associated with desiccation tolerance

We observed considerable genetic variability among DGRP lines for desiccation tolerance using a total of 162 lines (Table S1). Line means are depicted in Fig. 5. Females survived longer than males under desiccating conditions, as expected. Using this among line variation in the standard mapping pipeline (Mackay et al. 2012), SNPs associated with desiccation tolerance mapped to or close to 36 genes (Table S2 and Table S3).

**Fig. 5:**
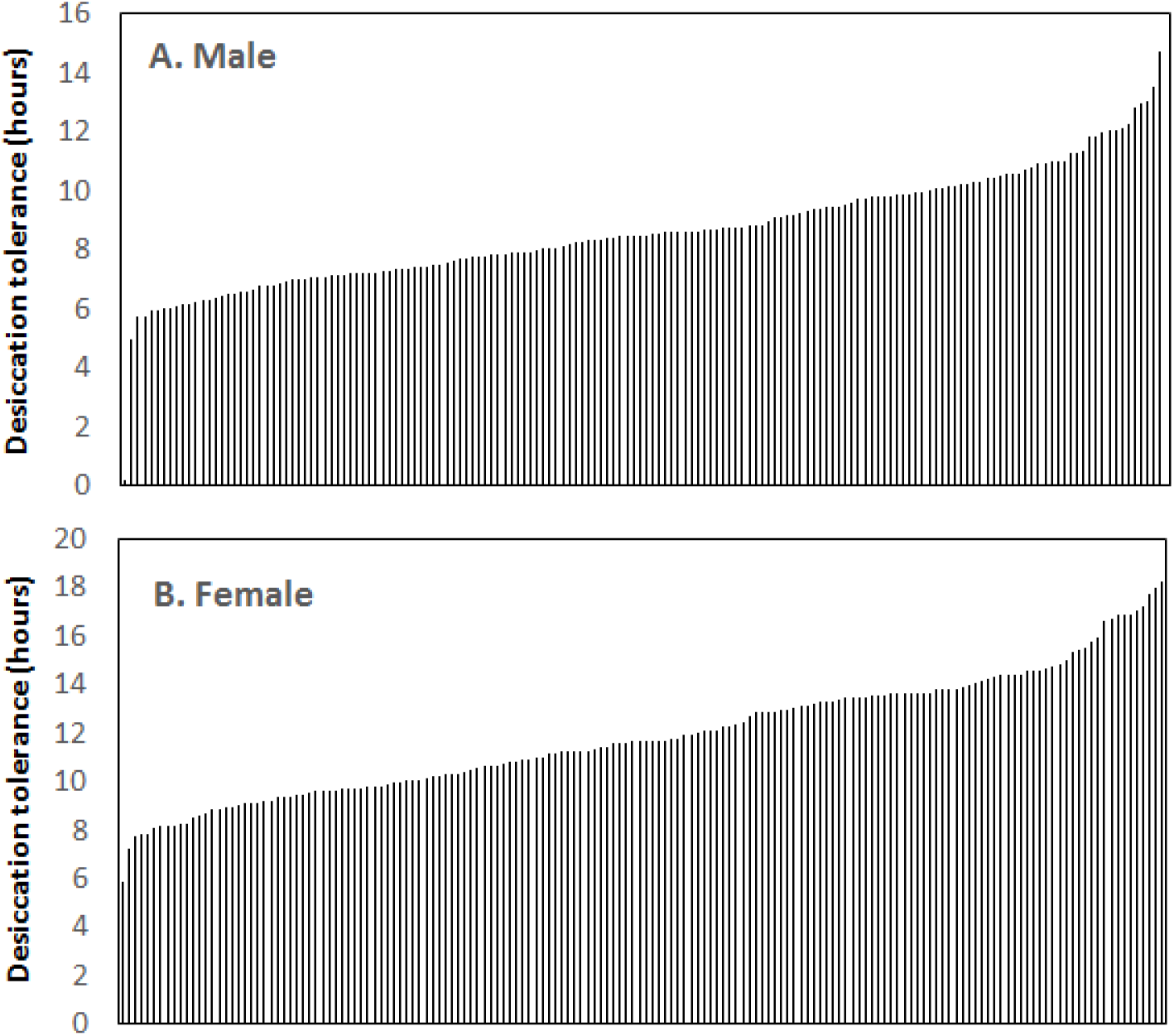
Status of desiccation tolerance (line means) in Drosophila Genetic Resource Panel. A total of 162 lines were considered for this association mapping. A considerable genetic variability in desiccation tolerance was observed across the lines. Males (A) survived shorter under desiccating conditions than females (B).

Among North American populations, genes associated with desiccation tolerance did not show increased signatures of spatially varying selection relative to the rest of the genome (Fig. 6; p=0.8), whereas they do among Australian populations (Fig. 6; p=0.02). The probability that *SUMSTAT*_*target*_ is greater than *SUMSTAT*_*control*_ was 0.74, indicating that genes associated with desiccation are not more differentiated among North American populations than expected relative to the rest of the genome.

**Fig. 6:**
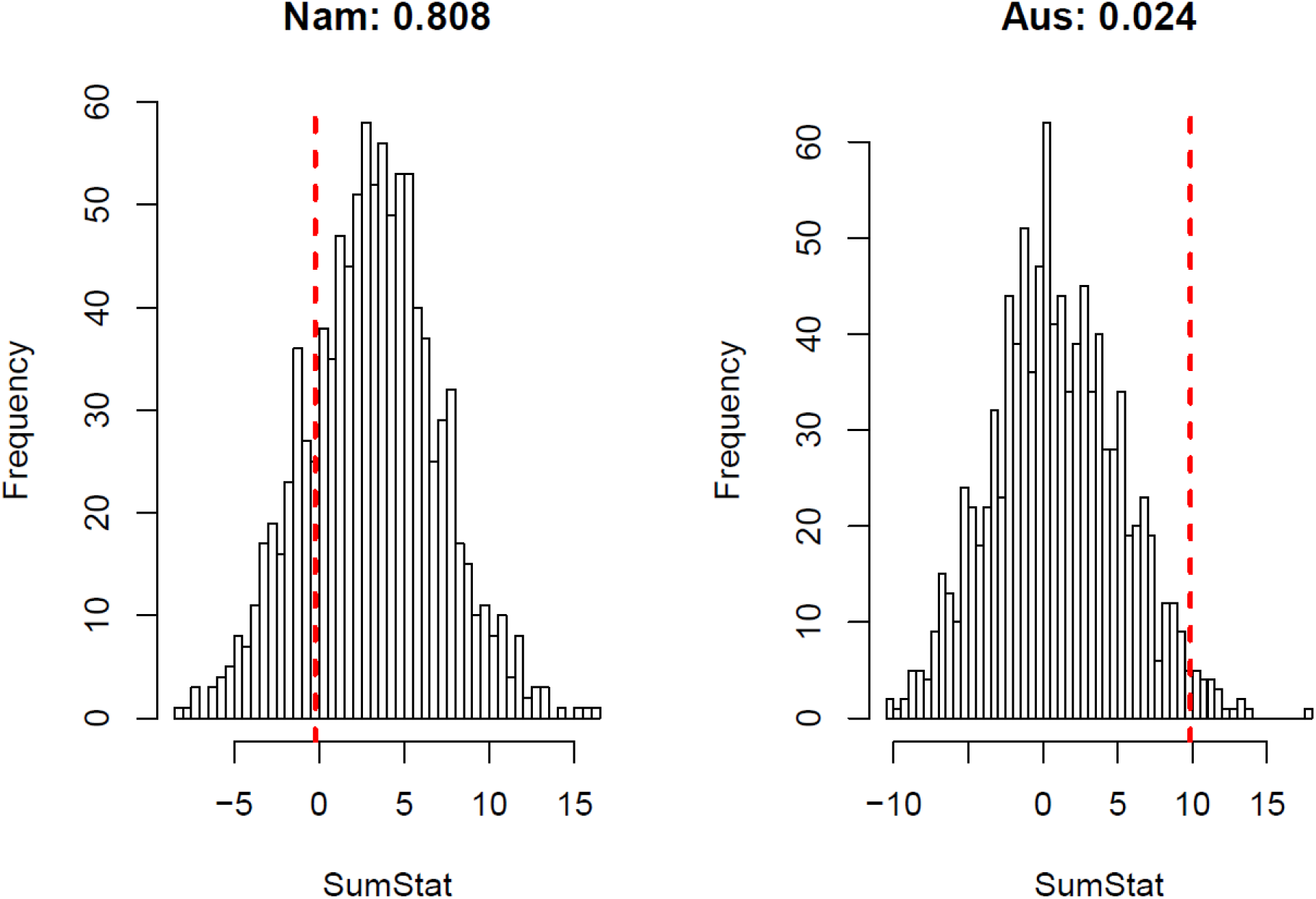
SumStat scores for 1000 sets of control genes (distribution) and observed SumStat score (dashed red line) for genes associated with desiccation tolerance among populations sampled along the east coast of North America (Nam) or Australia (Aus). See Materials and Methods for a description of the SumStat score.

## Discussion

Desiccation resistance in insects could involve one or more adaptive mechanisms, including (1) increases in total body water content and/or in haemolymph volume (Folk et al. 2001); (2) increased dehydration tolerance (i.e., tolerance of body water loss before death) (Telonis-Scott et al. 2006); (3) reduction in rate of water loss (Gibbs et al. 1997). In insects, higher surface area to volume ratio turn them susceptible to desiccation under drier climatic conditions. (which varied five-fold among 20 *Drosophila* species) and desiccation tolerance but not with rate of water loss. Based on these interspecific comparisons, it is generally predicted that body size can play a role in desiccation resistance. Along the coast of eastern Australia, no cline was observed for desiccation tolerance in *D. melanogaster* (Hoffmann et al. 2001), whereas along the Indian latitudes a robust cline has been observed (Karan et al. 1998). Interestingly, a cline for body size (e.g., thorax and wing length) has been observed on both continents (James et al. 1995; Bhan et al. 2014). In North America, body size also varies clinally and increases with increasing latitude (e.g., Coyne and Milstead 1987). Here, we observed a significant but shallow cline in desiccation tolerance at normal assay temperatures of 25˚C (Table S4). While body size may play a role in desiccation tolerance, the relationship between variation in body size and desiccation tolerance among *D. melanogaster* populations appears complex (Rajpurohit et al. 2016b, unpublished).

### Spatiotemporal patterns in desiccation tolerance

Our results, demonstrating a positive cline in desiccation tolerance for populations from the east coast of the U.S, are consistent with previous studies showing higher desiccation tolerance in temperate vs. tropical locales (Hoffmann & Harshman 1999). On the Indian subcontinent, this trend was clearly observed in multiple *Drosophila* species (Karan et al. 1998; Parkash et al. 2008; Rajpurohit & Nedved 2013), where parallel clines for this trait have been observed in several *Drosophila* species. On the Indian subcontinent higher latitudes in the north are characterized by lower temperature and lower humidity during winter, whereas low latitude, southern locations are warm and humid for most of the year. A meta-analysis approach concluded that T_cv_ (coefficient of variance in temperature) was a major climatic component to support the observed parallel clines for desiccation tolerance in several *Drosophila* species on the Indian subcontinent (Rajpurohit et al. 2013a). Along the east coast of the U.S., more temperate populations experience greater environmental fluctuations associated with seasonality and harsher winter conditions. As the temperate winter is generally associated with a highly desiccating environment, we predicted that populations collected in the spring would be characterized by relative increases in desiccation tolerance, similar to patterns observed for other stress-related traits (Behrman et al. 2015). However, we did not observe any differences in desiccation tolerance between early and late season collections of *D. melanogaster* from three temperate populations spanning 38 – 42˚N latitude. Similarly, we did not observe any differentiation in water loss rates in response to experimental evolution to seasonality in our field experiment. Humidity differences between fall vs spring seasons are not significant for any of these collection sites (e.g., Fig. S2), and this may be associated with the absence of seasonal variation in desiccation tolerance.

### Climatic associations

The widespread occurrence of latitudinal clines for many fitness related phenotypes may be related to the regularity of climatic changes, and associated parameters, with latitude. We analyzed climatic data along a south-north transect for the six sampled localities ranging from Florida (25.48 °N Latitude) to Maine (42.26 °N Latitude). Latitude was negatively correlated with winter temperature. We also observed that the amplitude of thermal seasonal variation, estimated by the between-month coefficient of variation (CV), increased with increasing with latitude, from 3.3% in Florida up to 29.7% in Maine (Schmidt et al. 2005).

It is generally assumed that desiccation tolerance is selected during hot and dry conditions, as heat and desiccation stresses generally co-occur (Hoffmann and Parsons 1991). In natural habitats on the U.S. east coast, however, such is not the case. Variation in desiccation tolerance is most strongly associated with minimum temperature of the geographic origin of our collections, suggesting that desiccation tolerance may be favored in environments characterized by cold and dry conditions (Leather et al. 1993). Cold and desiccation tolerance may also exhibit correlated responses (e.g., Bubliy and Loeschcke 2005; MacMillan et al. 2009). Thus, it remains unclear whether the latitudinal patterns we observed are driven primarily by selection directly on desiccation tolerance or may reflect an indirect response due to selection on a correlated trait.

### Thermal plasticity

Populations of *D. melanogaster* grown at lower temperatures slightly increased their desiccation tolerance. The difference in desiccation tolerance hours was in the direction of 18>25>29 °C (see slope comparison in Table S4). A recent study on the cold-adapted *D. nepalensis* from the western Himalayas found that flies grown at 15 °C show twofold higher body size, greater melanization, higher desiccation resistance, hemolymph and carbohydrate content as compared to flies reared at 25 °C (Parkash et al. 2014). There is a strong possibility that *D. melanogaster* populations growing at lower temperatures may also exhibit these plastic responses that could subsequently affect desiccation tolerance. However, we also observed that variation among populations became exacerbated at lower temperatures, and was most distinct following exposure to dormancy inducing conditions. Thus, our data also suggest that patterns of plasticity in *D. melanogaster* may vary predictably among natural populations and habitats.

### Geographic variation in metabolic rate

Desiccation resistance in *Drosophila* is associated with reduced water loss rates under both natural (Kalra et al. 2014) and laboratory conditions (reviewed by Hoffmann and Harshman 1999). In contrast, evidence for other potential adaptive mechanisms is more equivocal. Higher body water content was reported for resistant populations in some studies (Gibbs et al. 1997; Chippindale et al. 1998; Folk et al. 2001; Gefen et al. 2006), but not in others (Hoffmann and Parsons 1993). The ability to tolerate dehydration has also reported to vary between desiccation-selected populations and their controls in one study (Telonis-Scott et al. 2006), but not another (Gibbs et al. 1997). However, this discrepancy could simply reflect an inconsistency in the use of the term dehydration tolerance (Gibbs and Gefen 2009).

We found no evidence for variation in water loss rates that could explain the clinal variation in desiccation tolerance. Water-loss rates of the northernmost (Maine) and southernmost (Florida) populations did not differ in either males or females. We also recorded similar metabolic rates, expressed as CO_2_ emission rates 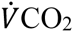 for females from the two populations, but higher values for severely desiccated Florida male flies suggest significant differences in the metabolic response of these populations to prolonged exposure to desiccating conditions. This is in agreement with both intraspecific (Hoffmann and Parsons 1993; Gefen and Gibbs 2009) and interspecific (Gibbs et al. 2003) reports which showed that desiccation resistance in *Drosophila* is correlated with reduced activity under stressful conditions; these results are also consistent with pronounced differences in central metabolism between populations at the geographic extremes of the U.S. east coast (e.g., Verrelli and Eanes 2001; Flowers et al. 2007; Lavington et al. 2014). It should be noted that increasing 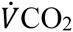 values could reflect a switch to carbohydrate catabolism under desiccation stress (Marron et al. 2003), independent of changing metabolic rates. However, we did not observe a similar response in females. Instead, the significant decrease in 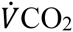 in the more resistant Maine females as they settled to the dry metabolic chamber environment, and absence of this response for Florida females, suggests a difference in behavioral response. Both males and females in these populations do exhibit very distinct behavior in response to thermal variation as well (Rajpurohit and Schmidt 2016).

Variation in activity patterns under stressful conditions is likely to result in correlated differences in respiratory water losses. The similar WLRs reported here for Florida and Maine flies can be explained by the considerably higher relative importance of cuticular water loss in insects (Chown 2002), and may suggest that flies across the experimental populations do not vary in their cuticular resistance to water loss. Nevertheless, while results in this study do not confirm an effect of activity level on WLRs and thus on desiccation-resistance, they could well reflect how stressful the exposure to experimental desiccation is to flies from the respective populations. If the more susceptible Florida flies have lower body water contents when hydrated compared with the more resistant Maine flies, then at similar WLRs the former would approach the minimum tolerable hydration state earlier, which could elicit an increase in activity levels as a result of attempts to seek more favorable conditions. In addition, a delayed escape response in the more resistant flies could indicate higher dehydration tolerance that would trigger an escape response at lower body water content.

### Ecological genetics of desiccation tolerance

Relative to other fitness-associated traits (e.g., body size, Coyne and Milstead 1987), reproductive dormancy (Schmidt et al. 2005), cuticular hydrocarbons (Rajpurohit et al. 2016c unpublished), thermal preference (Rajpurohit and Schmidt 2016) and body pigmentation (Rajpurohit et al. 2016a unpublished), we observed a very shallow cline for desiccation tolerance across the sampled latitudinal gradient in eastern North America. Our results also demonstrated pronounced patterns of plasticity in response to temperature, both in terms of developmental plasticity as well as adult acclimation and subsequent response. However, the observed patterns suggest that spatially varying selection may be less pronounced on this trait, both in comparison to other traits in North American populations as well as to desiccation tolerance in Australian (Telonis-Scott et al. 2006) and Indian (Karan et al. 1998) populations. Work done by Telonis-Scott et al. (2015), in comparison to our GWAS, further suggests the genetic basis of desiccation tolerance may be more robust and polygenic in Australian populations relative to North America. Similarly, our analysis of clinal enrichment demonstrated that genes associated with desiccation tolerance are enriched for clinality in Australia but not in North America. Differences among the continents may reflect differential local adaptation and the role of desiccation tolerance in affecting fitness, variation in colonization history and associated demography (Kao et al. 2015; Bergland et al. 2016), or a combination of the two.

### Conclusion

We observed a shallow cline for desiccation tolerance in populations sampled along the latitudinal gradient in the eastern U.S.; these patterns of variation among populations exhibited both developmental and adult plasticity, suggesting that further analysis of desiccation tolerance should be examined under a range of environmental conditions. Climatic analysis of this cline indicated that observed patterns of desiccation tolerance were most strongly associated with lower temperature conditions, suggesting that selection on this trait in temperate populations may be associated with response to desiccating conditions that co-occur with exposure to reduced temperatures. GWAS analysis using the DGRP panel of inbred lines identified 36 genes associated with desiccation tolerance in North American populations, providing a wealth of candidates for subsequent functional analysis and investigation. These genes were not enriched for signatures of clinality in North American populations but were in Australian populations, further suggesting differential dynamics of this trait in various habitats of this cosmopolitan genetic model organism.

## Data Accessibility

Raw data will be archived at the Dryad digital repository.

## Acknowledgements

We would like to thank Kelly Dyer for providing isofemale lines collected from Athens, GA and Emily Behrman for providing seasonal collections from Media, PA. Research in the Schmidt Lab & Gibbs Lab was supported through the National Institute of Health grant R01GM100366 and National Science Foundation (EnGen-0723930) respectively. This work was also supported through a grant from the Peachey Environmental Biology Fund to SR. We are particularly thankful to undergraduate research assistants in the Gibbs Lab for their 24-hour help in phenotyping DGRP lines.

## Supporting Figures

**Fig. S1**: qq-plot of genome wide association analysis of intersex averaged desiccation tolerance. Observed p-values are plotted along the y-axis and expected p-values along the x-axis. The solid red line depicts the 1:1 line.

**Fig S2**: 3D Surface plot (A) of T_ave_ and RH_ave_ along the latitudes of east coast of U.S.A. RH_ave_ = Distance weighted least square. Geographical clines in fitness traits and their interactions with climatic variables are well known. To explore these interactions in this study we planned to study the relationship between desiccation tolerance and climatic variables (temperature and relative humidity) of the origin of sites of populations. Temperature and relative humidity data for sample collection sites averaged over 30 years (from 1980 to 2010). Seasonal variations (Spring vs. Fall) in relative humidity of the origin of sites of population collections are also presented (B). All the historical data of temperature and humidity were downloaded from the National Climatic Data Center (NOAA; http://www.ncdc.noaa.gov/).

